# Musculotendon Parameters in Lower Limb Models: Simplifications, Uncertainties, and Muscle Force Estimation Sensitivity

**DOI:** 10.1101/2022.11.04.515177

**Authors:** Ziyu Chen, David W. Franklin

## Abstract

**Objective:** Musculotendon parameters are key factors in the Hill-type muscle contraction dynamics, determining the muscle force estimation accuracy of a musculoskeletal model. Their values are mostly derived from muscle architecture datasets, whose emergence has been a major impetus for model development. However, it is often not clear if such parameter update indeed improves simulation accuracy. Our goal is to explain to model users in which way and how accurate these parameters are derived, and to what extent errors in parameter values might influence force estimation.

**Methods:** We examine in detail the derivation of musculotendon parameters in six muscle architecture datasets and four prominent OpenSim models of the lower limb, and then identify simplifications which could add uncertainties to the derived parameter values. Finally, we analyze the sensitivity of muscle force estimation to these parameters both numerically and analytically.

**Results:** Nine typical simplifications in parameter derivation are identified. Partial derivatives of the Hill-type contraction dynamics are derived. Tendon slack length is determined as the musculotendon parameter that muscle force estimation is most sensitive to, whereas pennation angle is the least impactful.

**Conclusion:** Anatomical measurements alone are not enough to calibrate musculotendon parameters, and the improvement on muscle force estimation accuracy will be limited if the source muscle architecture datasets are the only main update.

**Significance:** Model users may check if a dataset or model is free of concerning factors for their research or application requirements. The derived partial derivatives may be used as gradients for musculotendon parameter calibration. For model development, we demonstrate that it is more promising to focus on other model parameters or components and seek alternative strategies to further increase simulation accuracy.

## I. Introduction

Human movement is actuated by muscles, and perhaps the fact that *kinesiology* (the study of human movement) is often referred to as human *kinetics* (the study of forces) highlights the importance of musculoskeletal forces in research. Nowadays, medical and kinesiology professionals are increasingly interested in the vast information hidden in kinetic data. For example, clinicians require for diagnosis and surgery the knowledge of the specific muscles and biomechanical properties responsible for a movement dysfunction. Athletic and rehabilitation trainers benefit from understanding how task performance can be improved or how joint load can be alleviated by exercising certain muscle groups. In the field of motor control, the timing and intensity of muscle activities provide valuable insight into the regulation of movement in the everchanging environment.

Nevertheless, by its very nature, human movement research has limited accessibility to the kinetics of the musculoskeletal system. Direct in vivo measurement of human joint and muscle forces is arduous along with ethical concerns [1], [2]. Thus, biomechanical investigations are often confined to kinematic analysis, with very few direct kinetic measurements available. Instead, most rely on indirect techniques such as dynamometry and electromyography (EMG).

While a dynamometer measures isometric or isokinetic joint moments of force [3], it is not compatible with most motor tasks due to its design. To estimate muscle forces in dynamic motion, EMG is frequently used as a measure of the intensity of muscle activation. However, apart from the difficulty to obtain EMG from deep muscles, especially during movement, the correlation of EMG and muscle force is limited [4], particularly in concentric and eccentric contractions.

The pursuit of accurate kinetic estimation has led to extensive development of musculoskeletal modeling in biomechanical research over the last few decades. A musculoskeletal model is a set of equations describing the musculoskeletal system [5] via mathematical connections between neural signals, muscle forces, joint moments, and skeletal motions. The kinetics and neuromuscular signals can either be inversely estimated from measurements of kinematics and external reaction forces, or they can be predicted in specific tasks given a set of objective functions [6]–[14].

Most musculoskeletal models use the Hill-type muscle model, whose contraction dynamics are mainly determined by four musculotendon parameters: optimal fiber length, maximal isometric force, pennation angle, and tendon slack length. These parameters reflect real physiological quantities directly derivable from the musculoskeletal system, and the parameter values themselves are worthy of attention during the investigation of human anatomy. This feature distinguishes musculoskeletal modeling from other model-based kinetic estimation techniques. For instance, it is recently demonstrated how a neural network can accurately estimate muscle and joint forces from data as simple as ground reaction forces, outperforming any manually designed musculoskeletal models [15]. In this case, the neural network serves only for calculation, while the physiological representation of each node is unknown, making it difficult to gain any new perspectives on the musculoskeletal system. In contrast, musculoskeletal models provide a more straightforward relation between the simulation results and the musculoskeletal architectures, allowing direct biomechanical insight [16]–[18]. In short, musculotendon parameters not only are a key to kinetic analysis, but also play a crucial part in how we understand the musculoskeletal system.

Human measurement is a major source for musculotendon parameters, and the emergence of new muscle architecture datasets often urges the development of new models, especially for lower limb models. Initial data collected by Wickiewicz et al. [19] on three elderly cadavers, and Friederich and Brand [20] on two elderly cadavers served as the gold standard references for research in muscle biomechanics, including Delp’s [21] model. Because of the scarcity of available data, new measurements from a single elderly cadaver [22] drove the development of a new model [23]. However, soon after Ward et al. [24] published their dataset based on as many as 21 elderly cadavers, Arnold et al. [25] developed their lower limb model with this parameter update as a major feature. To avoid the effect of old age on muscles, Handsfield et al. [26] took the measurement in vivo using MRI on 24 young subjects, which Rajagopal et al. [27] employed in their model. Recently, Charles et al. [28] provided a similar MRI-based dataset that attempted in vivo measurement of fiber length, which could potentially motivate yet another upgrade of lower limb models.

Although it seems logical to drive model development with up-to-date muscle architecture data, the concerns are two-fold. First, cadaveric and MRI data are not equivalent to musculotendon parameters; nor is the validity of each datum or parameter guaranteed. Simplifications in experimental measurements are less of a focus in the modeling phase where the values may receive more attention than the methods, and this issue is further compounded in calculating musculotendon parameters from the raw data. This means that earlier simplifications are often inherited into new models, potentially affecting simulation accuracy without being explicitly considered or presented.

This first concern is increasingly evident as many researchers, both inside and outside academia, benefit from the advancement of modeling software. In particular, OpenSim [29] is an open-source software created by modeling professionals which provides easy access to dynamic simulations and allows convenient development of musculoskeletal models, and it is one of the pre-eminent platforms in musculoskeletal modeling. Nevertheless, precisely because of this level of prominence, many users overlook the potential issues with the original derivation of the parameters within OpenSim models. Many of these simplifications were necessary to deal with problems unclear at the time of modeling, and may not be solved in the present, but can contribute to inaccuracies in simulation results.

The second concern is to what degree do model simulations benefit from higher accuracy in the musculotendon parameters, and whether there is a limit to such improvement. Musculoskeletal models and their components, including the Hill-type muscle model, are highly non-linear, which means that improvements in the accuracy of partial parameters do not necessarily lead to improvements of the overall simulation results. More particularly, there may be specific parameters that are much more critical for model calibration.

In this work, we discuss how musculotendon parameters in lower limb models were derived and how some of the simplifications in this process produce uncertainties in parameters that can affect force estimation. We extend this by demonstrating how muscle force estimation is affected by errors in each parameter, both numerically and analytically.

## II. Definition

Here, we examine the derivation of musculotendon parameters in six muscle architecture datasets and four prominent OpenSim musculoskeletal models of the lower limb (Table I). We attempt to distinguish between decisions about parameters or parameter values that are *simplified* from those based on evidence. Specifically, we use the term *simplifications* to refer to decisions without traceable references in support of the specific cases. For example, it has been well studied that maximal isometric force is related to physiological cross-sectional area (PCSA) of muscle [18], [30], [31]. Therefore, it is not considered a simplification, but rather a reasonable decision, for modelers to estimate maximal isometric force from the product of PCSA and specific tension. However, there is no supporting evidence that all PCSAs from the same dataset should be scaled with a uniform specific tension. This is a simplification and needs further discussion. Finally, by *derivation*, we refer to both the measurement of muscle architecture data and the calculation of musculotendon parameters from the data.

**TABLE I.**
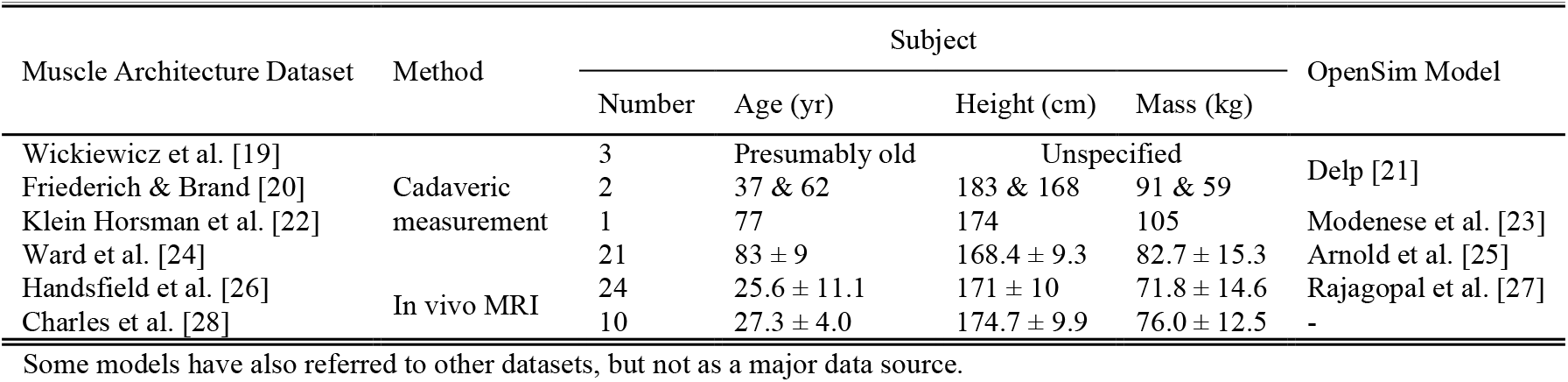
Major lower limb muscle architecture datasets and OpenSim models

Importantly, given the limited information available for muscle architectures and kinetics, we cannot determine which parameter value is inherently correct for muscle force estimation. Instead, by *uncertainty*, we refer to the difference in value between how the parameter is supposed to be derived by definition and how it was actually derived. It is beyond the scope of this paper to specify all musculotendon parameter uncertainties in each dataset and model. Instead, we focus on providing examples of representative cases where the differences may be large due to simplifications. The goal is to highlight the way in which these simplifications might lead to uncertainties, and to show the extent to which these could influence muscle force estimations. Model users can then evaluate if a particular parameter should be scrutinized for the requirements of their specific scientific questions. For example, should parameters derived from elderly cadavers be used for simulations of young athletes.

In order to achieve ideal simulation of the musculoskeletal system, we require accurate descriptions of all aspects of a musculoskeletal model, including:

- *Skeletal geometry*. Describes skeletal mass distribution and the joint motions during a task.
- *Musculoskeletal geometry*. Describes the muscle length and moment arm in a given joint position.
- *Multibody dynamics*. Describes the moments of force needed to generate given joint motions and external forces, or the joint motions and external forces generated from given moments of force.
- *Neural control principle*. Describes the force each muscle generates when a given joint moment is needed, typically referred to as the *force distribution problem* [32], [33].
- *Contraction dynamics*. With fiber length and velocity given, describes the muscle activation needed to generate a given force, or the force generated from a given muscle activation.
- *Activation dynamics*. Describes the neural excitation needed to generate a given muscle activation, or the activation generated from a given neural excitation.

In this paper, we focus only on contraction dynamics. More specifically, we focus on the equation of the Hill-type muscle contraction dynamics:

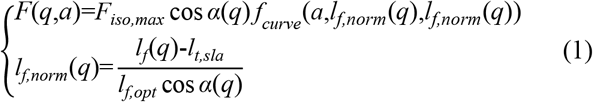

where *F*(*q,a*) is the muscle force developed at some joint position *q* and muscle activation level *a. l*_*f*_(*q*) denotes the fiber length at *q*: *l*_*f,norm*_(*q*) is its normalized value, and 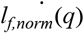 is the normalized fiber velocity. *F*_*iso,max*_ is maximal isometric force, *l*_*f,opt*_ is optimal fiber length, *α*(*q*) is the pennation angle at *q, l*_*t,sla*_ is tendon slack length, and the function *f*_*curve*_(*·*) defines the four Hill-type characteristic curves, including the active and passive muscle force–length curve, the muscle force–velocity curve, and the tendon force–length curve.

In later sections, the term *force estimation* is implicitly referred to as in an approximate calculation of muscle force with Eq. 1. Hence it is also implied that the maximal accuracy of force estimation does not go beyond the limit set by the Hill-type muscle model itself. That is, with all simplifications eliminated and the best possible parameters derived for Eq. 1, estimated force could still contain errors, but these would be due to biomechanical mechanisms undescribed by the Hill-type muscle model.

## III. Derivation of Musculotendon Parameters

This section covers the four musculotendon parameters in the order of optimal fiber length, maximal isometric force, pennation angle, and tendon slack length. While they are key factors in Eq. 1, it should not be neglected that the parameters shaping the characteristic curves in a Hill-type model also play a role in muscle contraction dynamics. In Millard et al.’s [34] muscle model for example, the passive force–length curve can be modified by defining the strain where passive force begins to develop (0 by default) and reaches maximal isometric value (0.7 by default). Although the Hill-type characteristic curves are almost always left in the default configuration, they can be differently shaped across muscles [35], [36]. In addition, many studies suggest that the fiber length at which passive fiber force develops does not necessarily coincide with optimal fiber length, as often simplified in modeling [35], [37], [38]. Keep in mind that force curves are imposed under this default configuration only because they are more complicated to measure than the typical musculotendon parameters. Once there are available data, these parameters should be tuned.

### A. Optimal Fiber Length

Optimal fiber length is “the length at which maximal force can be produced.” [39, p. 34] This is both a biomechanical concept and an anatomical one. Yet neither cadaveric nor imaging measurements are possible while monitoring active individual muscle force, so the raw fiber length cannot be assumed as the optimal length. In addition, fibers are not without passive loading in the fixation position, where cadavers are seemingly *at rest*. Even when dissected, sarcomere length varies across muscles [24], indicating non-uniform deformation.

There are difficulties that can arise in the measurement and conversion of the raw fiber length to optimal fiber length. For example, Wickiewicz et al. [19] measured both fiber and sarcomere lengths for each muscle, and normalized the fiber length to a sarcomere length of 2.2 μm in PCSA calculation, which theoretically eliminates the influence of unknown muscle lengthening and shortening [40]. Delp rescaled the fiber length reported by Wickiewicz et al. [19] with a sarcomere length ratio of 2.8 to 2.2, with 2.8 μm being “the optimal sarcomere length” for it is “the length at which a fiber develops peak force based on the sliding filament theory of muscle contraction.” [21, p. 6] This approach would infuse biomechanical meaning into architecture data, making them theoretically optimal (*optimal at best*; Fig. 1, middle). However, although Wickiewicz et al. [19] emphasized this *optimal scaling* approach, it appears that scaling was exclusively conducted in PCSA calculation, meaning that the fiber length reported in their table is in fact unscaled raw fiber length (as can be confirmed by muscle volume calculation). We suggest that the rescaling of the muscle lengths was therefore ineffective, as the raw fiber length, rather than optimal fiber length, was scaled [21].

**Fig. 1.**
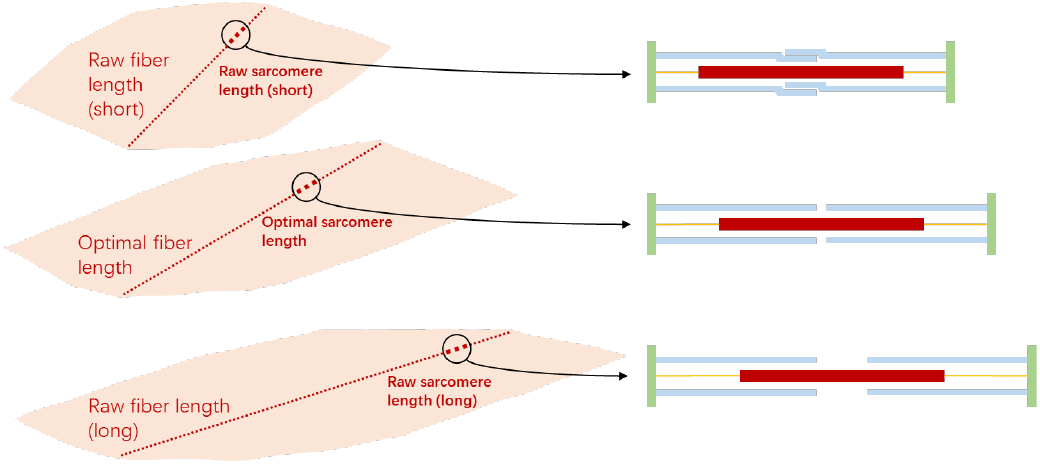
The relation between fiber and sarcomere length. A muscle fiber is consisted of sarcomeres connected in series, such that the lengths of the muscle fiber and sarcomere are positively correlated. A sarcomere is consisted of the myosin (red), actin (blue), and titin (yellow) filaments as well as Z membranes (green), and its length is measured as the distance between Z-membranes. A contractile force is generated when crossbridges are formed as a link between the myosin and actin filaments. Top: When the sarcomere is short enough for actin filaments to overlap with each other, fewer crossbridges can be formed, and less force can be generated. Middle: When the sarcomere is at a length where the myosin filament is fully covered by non-overlapping actin filaments, the most crossbridges can be formed, and the most force can be generated; this length is hence optimal. Bottom: When the sarcomere is short enough for the myosin filament to be less covered by actin filaments, fewer crossbridges can be formed, and less force can be generated.

For muscles not reported by Wickiewicz et al. [19], data from Friederich and Brand [20] were used by Delp [21]. The lengths were taken in the fixation or resting position with no scaling of any kind, which means that a simplification that fiber optimality occurs in the resting position is made (*optimal at rest*; Fig. 1, top & bottom) as there is no alternative solution. Care must be taken when looking at simulation results of these muscles.

Ward et al. [24], Klein Horsman et al. [22], and Handsfield et al. [26] adopted the same scaling approach but with a different optimal sarcomere length of 2.7 μm. This particular number comes from the study of Lieber et al. [41], where they estimated an optimal human sarcomere length of 2.60–2.80 μm, agreeing well with the ranges reported by Walker and Schrodt [42] (2.64–2.81 μm) as well as Gollapudi and Lin [37] (2.54–2.78 μm). If other values within these ranges are chosen, there will be slight differences in the optimal fiber length estimates.

Nonetheless, the optimal scaling approach simplifies sarcomere length to be uniform within each muscle, as well as the optimal sarcomere length to be constant across different muscles. Recently, Moo et al. [43] has shown that the sarcomere length in mouse tibialis anterior may differ up to 0.3 μm (nearly 15%) when measured in distal and proximal sites. Hessel et al. [44] reported a difference of 0.1 μm (about 5%) between the optimal sarcomere length of mouse soleus and extensor digitorum longus. This evidence indicates that sarcomere length uncertainties in previous cadaveric measurements might be larger than anticipated, and further uncertainty could be created when scaling with a constant optimal sarcomere length.

In spite of such problems in cadaveric measurement and optimal scaling, in vivo MRI does no better job in obtaining precise length data. Charles and colleagues measured fiber lengths with diffusion tensor imaging and concluded an accuracy of 1±7 mm [28], [45], but it is critical to keep in mind that the 1-mm error comes from averaging errors in all muscles of all subjects. This simply indicates that the overall technique is not specifically biased, not that the actual fiber length estimates are accurate (for example, the mean of white noise is zero). A standard deviation of 7 mm is also not particularly low for muscle fiber measurement. The soleus fiber length in three cadavers was measured both directly and through MRI, with results of 56±10 mm and 78±21 mm respectively [28]. However, in addition to these large differences, the soleus in a related dataset [28] is listed with a fiber length of 146±32 mm, which disagrees with both data above as well as previous results [19], [20], [24]. This method must be further investigated before such datasets are used for musculoskeletal modeling.

### B. Maximal Isometric Force

Maximal isometric force is the force capacity of a muscle in the isometric condition and is most difficult to measure muscle by muscle. As mentioned above, it can be estimated from specific tension (*σ*_*iso,max*_) and PCSA (*A*_*PCS,opt*_):

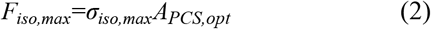

PCSA is the area of the cross section perpendicular to the fibers, usually calculated using Eq. 3.1 with specification of fiber optimality, as can be indicated from the notation. It is worth noticing that the concept of *projected* or *functional PCSA* is also often used [39, p. 53], where the cosine of pennation angle is additionally multiplied (Eq. 3.2), and the resultant value is therefore theoretically close to anatomical cross-sectional area (ACSA).

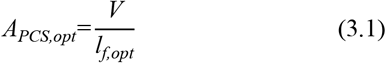

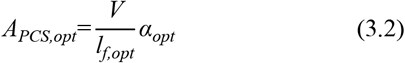

where *A*_*PCS,opt*_ denotes the PCSA when fiber length is at optimality. *V* is muscle volume, and *α*_*opt*_ is the pennation angle at optimal fiber length.

The preference over the two definitions varies. Delp [21, p. 5] defined PCSA in the unprojected fashion, but the value from Wickiewicz et al. [19] he used to calculate maximal isometric force was actually projected PCSA. Although Ward et al. [24] provided the projected PCSA, Arnold et al. [25] went through the trouble to recalculate PCSA with muscle volume and optimal fiber length from the same dataset.

Both choices are supported by studies, that the conventional PCSA, projected PCSA, and even ACSA all can be highly correlated to muscle force capacity [18], [30], [31]. However, considering that most lower limb muscles have pennation angles small enough for similar cosine values [19], [20], [24], [28], a decision between the two definitions cannot be made without testing its effects on muscles with large pennation angles, such as the gluteus and soleus.

More importantly, both two techniques involve optimal fiber length to calculate PCSA, so the accuracy of the PCSA estimate will be determined by that of the optimal fiber length measurement. Different starting estimates produce large variations in the estimates of PCSA. For example, as described previously, Charles et al. [28] determined the optimal fiber length of the soleus to be 146±32 mm, producing a projected PCSA of their young subjects of 32.3±10.4 cm^2^. However, this value is small compared to Ward et al.’s [24] estimation of 51.8±14.9 cm^2^ for their elderly specimens. The difference is further enhanced if the conventional PCSA is calculated, since pennation angle is measured to be 12°±2° in Charles et al. [28] and 28°±10° in Ward et al. [24]. It is very unlikely that the young subjects should have smaller and thinner soleus muscles, and in fact, the soleus volume in Charles et al. [28] is 77% larger than that in Ward et al. [24]. The problem is mostly that this volume gets divided by the large optimal fiber length, providing an estimate of PCSA in the younger subjects well below that estimated for elderly cadavers.

Similarly, the accuracy of pennation angle is also relevant in projected PCSA calculation. Although pennation angles of the lower limb muscles are generally small, meaning that the cosine values do not differ by much, we still need to consider one key point. Optimal fiber length is not the raw fiber length measured experimentally but is scaled with sarcomere length. However, this scaling process often uses the raw pennation angle measured experimentally. By using the scaled fiber length and raw pennation angle in projected PCSA calculation, pennation angle is held to remain unchanged when the fiber is stretched or shortened to the optimal length (Fig. 2).

**Fig. 2.**
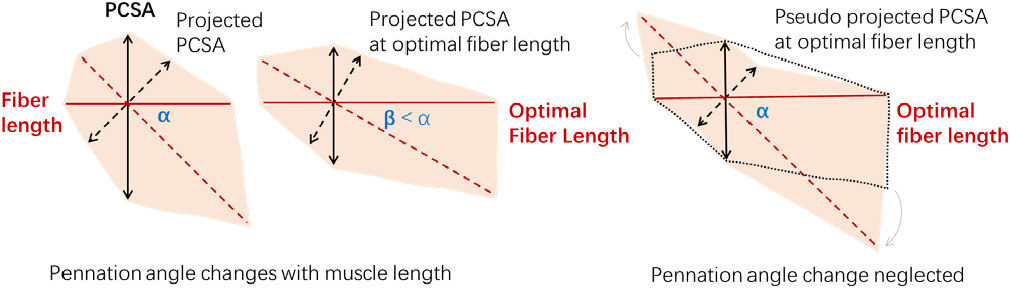
The PCSA change during muscle stretch when pennation angle changes along or remains unchanged. For convenience, fiber length (red solid line) is treated as muscle length (red dashed line) projected onto the pennation direction, and the projected PCSA (black dashed line) is hence PCSA (black solid line) projected onto the anatomical cross section. Since pennation angle decreases as the muscle gets stretched, the fiber- to muscle-length ratio increases, and the conventional to projected PCSA ratio decreases. This is not the case in the hypothetical situation where pennation angle stays unchanged. It would be similar to tilting and further stretching the muscle, and since the volume of muscle is constant, the extended area on both ends must come from the anatomical cross section, decreasing the projected PCSA.

The consequence of this simplification may not be trivial. Again, we will take the soleus as an example. With a sarcomere length of 2.12 μm measured by Ward et al. [24], it needs to be elongated nearly 30% to reach the proposed optimal sarcomere length of 2.70 μm. According to the data from Kawakami et al. [46] and Maganaris [47], such elongation is accompanied by a decrease of about 10% in the cosine value of the soleus pennation angle, equivalent to a 10% increase in the projected PCSA compared to a calculation without such consideration. This may explain why Arnold et al. [25] did not directly use the projected PCSA from Ward et al. [24].

A similar simplification occurred in Handsfield et al. [26] and Rajagopal et al. [27] when calculating the conventional PCSA. The muscle volume measured in vivo from young subjects [26] was divided by the optimal fiber length from elderly cadavers [24]. In Handsfield et al.’s [26] case, the value was further normalized with the muscle length ratio of the two datasets, but the problem remains fundamentally the same. Here, the fiber length is held to remain unchanged as age increases, yet there is evidence of significant fiber length reduction in aged mice and rats [48]–[50]. It is therefore possible that the mixed use of two datasets with differently aged subjects overestimates PCSA. Since there is not yet a reliable way to measure fiber length in vivo, the calculation of PCSA from the recent MRI-measured muscle volume will inevitably require the optimal fiber length from cadaveric measurement, but the potential overestimation induced by this *inherited fiber length* simplification should be carefully considered depending on the research requirements.

Importantly, the structure of Eq. 2 highlights the issue that an accurate PCSA still requires an appropriate value of specific tension in order to estimate maximal isometric force. In the early days, only PCSAs from elderly cadavers were available, so an atypical specific tension was needed to scale the force production capacity for a generic model of young adults. The value of 61 N/cm^2^ was first used by Delp in his PhD thesis, because it met the need to “match the moment curves measured on young subjects.” [21, p. 34] Although not experimentally derived, this value is in some sense optimization-derived, and proves to be a reasonable choice for muscle volume data from elderly cadavers, as was later qualitatively validated by Arnold et al. [25]. However, this value should not be used as a gold standard. As stated earlier, Delp [21] rescaled the fiber length from Wickiewicz et al. [19] with a factor of 2.8 to 2.2 to ensure theoretical optimality, but maximal isometric force was calculated using the PCSA from Wickiewicz et al. [19], which was based on the un-rescaled short fiber length. Therefore, Delp’s specific tension value was based on shorter and wider muscles. If instead, this value was based on the PCSA calibrated with the rescaled optimal fiber length, it would be 78 N/cm^2^. Importantly, this correction does not conflict with the validation by Arnold et al. [25] using 61 N/cm^2^, since they ran a qualitative comparison, whose results remain satisfactory even if scaled up by a fraction.

In later work, Rajagopal et al. [27] directly inherited Delp’s specific tension (determined as a scaling factor to convert elderly cadaver data to match younger experimental results) but based on the data collected in vivo from young subjects. This is a simplification of *inherited specific tension*, and we argue that the use of this high specific tension in such a case would overestimate maximal isometric force. Indeed, their model evaluation demonstrated much higher joint moments compared to experiment results, exactly against the original support of this value of specific tension [21, p. 34]. Although they noted that there was the need to increase muscle force capacity to simulate dynamic motions such as running, the question remains whether all muscles need to be scaled with a high specific tension, even for sake of simulation.

This brings us back to the problem pointed out earlier, that specific tension is simplified to be uniform across all muscles and muscle fiber types. The soleus, for instance, consists predominantly of slow-twitch fibers, which could have a lower specific tension than fast-twitch fibers [51], [52]. Kawakami et al. [53] and Buchanan [54] reported different specific tensions between human elbow flexors and extensors, and Fukunaga et al. [55] reported a two-fold difference in specific tensions between ankle plantarflexors and dorsiflexors. A comparison of these studies also suggests that upper limb muscles have much larger specific tension than lower limb muscles. Such factors raise questions about whether Rajagopal et al. [27] can use Buchanan’s [54] estimation of 100 N/cm^2^ in elbow flexors in support of the 60 N/cm^2^ they selected for lower limb muscles. Similarly, variations exist even for muscles in the same functional group. Javidi et al. [56] directly measured muscle force in kangaroo rats and found a 25% larger specific tension in gastrocnemius than plantaris. If accurate maximal isometric forces are regarded as a key feature in modeling, then modelers should invest as much, if not more, time in choosing specific tensions as they did for PCSAs.

It is also worth pointing out that we should be careful about directly inheriting specific tensions reported in literature. Specific tension is “maximal muscle force per unit of the muscle PCSA,” [39, p. 159] but it is critical to note that the force of a fully activated muscle also depends on its length and velocity, which must be specified. The large range of reported experimental values partially arises through the result of different muscle contraction states rather than the physiological heterogeneity in the muscles themselves. For modelling purposes, the specific tension values should be the maximal values measured under isometric conditions. For example, Kawakami et al. [53] estimated isokinetic tensions for different velocities, but only the ones measured without motion would be useful for parameter derivation. Similarly, Fukunaga et al. [55] provide a variety of estimates of isometric tensions in different joint angles, only the largest of which should be used.

Due to the scarcity of specific tension data, often values from literature must be adopted, despite undesirable mismatch of the subjects or muscles. Modenese et al. [23] decided on 37 N/cm^2^ for the lower limb muscle specific tension based on two studies. The first, by Haxton [57], is restricted by past measurement techniques, where PCSA was approximated via calf circumference and more simplifications were made to measure force. The second reference is the measurement by Weijs and Hillen [58], [59] on the human jaw muscles, which are intuitively much stronger than lower limb muscles. In Delp’s [21] model, for muscles whose data were obtained from a relatively young cadaver by Friederich and Brand [20], he used a specific tension of 25 N/cm^2^, referring to Spector et al.’s [60] measurement of 23 N/cm^2^. This value was obtained from the triceps surae, which can be suitable for other lower limb muscles, but the subjects were cats. Nevertheless, this is not to say that any of these choices of specific tensions are inappropriate (in fact, kinetic validations suggest these values may be in the correct range), but only to emphasize the necessity of re-evaluating dataset features when more data become available in literature.

### C. Pennation Angle

Pennation angle is measured between the direction of the muscle fiber and the direction of the tendon and determines how much force can be transmitted. In the previous subsection, a simplification of *fixed pennation angle* is pointed out, where not considering the variation of pennation angle with muscle length could induce error in projected PCSA calculation.

Essentially, PCSA and pennation angle are variables that change during motion, yet a Hill-type muscle model is defined by constant parameters. What modelers really need for musculotendon parameters are the architecture data measured when muscle exerts maximal active force. Without such measurements, care needs to be taken while processing the values to adjust them to the optimal states. Indeed, such adjustment explains the optimal scaling of fiber length, as well as the division of muscle volume by its own optimal fiber length in PCSA calculation. The same goes for pennation angle, which has to be measured at optimal fiber length or scaled; otherwise, it is a simplification of *optimal at rest*.

Unlike fiber length, pennation angle has no quasi-linear relation with sarcomere length, so its normalization is more complicated. Equation 4.1 describes the dynamic change of pennation angle assuming a constant muscle thickness [61]:

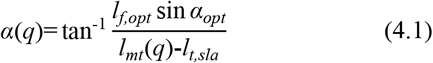

where *α*(*q*) and *l*_*mt*_(*q*) respectively denote pennation angle and muscle–tendon unit (MTU) length at some joint position *q*.

If sarcomere length is available from measurement, Eq. 4.1 can be transformed as:

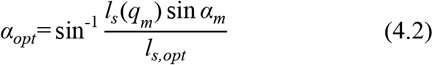

where *α*_*m*_ and *l*_*s*_(*q*_*m*_) are the measured pennation angle and sarcomere length, and *l*_*s,opt*_ is the optimal sarcomere length discussed in the previous section.

With either Eqs. 4.1 (requires definition of musculoskeletal geometry) or 4.2 (requires sarcomere measurement), the optimal scaling of pennation angle can be performed. Of all models in Table I, this is only accomplished in Rajagopal et al. [27], where the difference is evident for muscles with large pennation angles. For example, the optimal pennation angle of the soleus is corrected as 21.9° from the measured 28.3° [24], which is approximately 5% change in cosine value.

### D. Tendon Slack Length

By convention, tendon slack length refers to the length of the tendon where it starts to generate a restoring force to any change in its length, but it will be shown that a different concept of tendon slack length is being used in modeling. Due to the difficulty of accurately measuring tendon length, most datasets in Table I do not provide relevant values, and Arnold et al. [25] and Rajagopal et al. [27] used Eq. 5 to set tendon slack length:

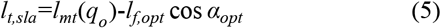

where *l*_*mt*_(*q*_*o*_) is the MTU length at a predetermined joint position *q*_*o*_.

The key point in this equation is MTU length, which varies depending on the joint position *q*_*o*_. Thus, setting the tendon slack length is equivalent to selecting the joint position for the fiber to be in its optimal length (Fig. 3). One of the biggest simplifications in parameter setting is that fiber optimality occurs in the joint position that measurement took place. In other words, the fixation or resting position (*q*_*meas*_) is the *optimal joint position* where maximal isometric force is compelled to appear, consequently determining how the force– length curve is expressed in each muscle.

**Fig. 3.**
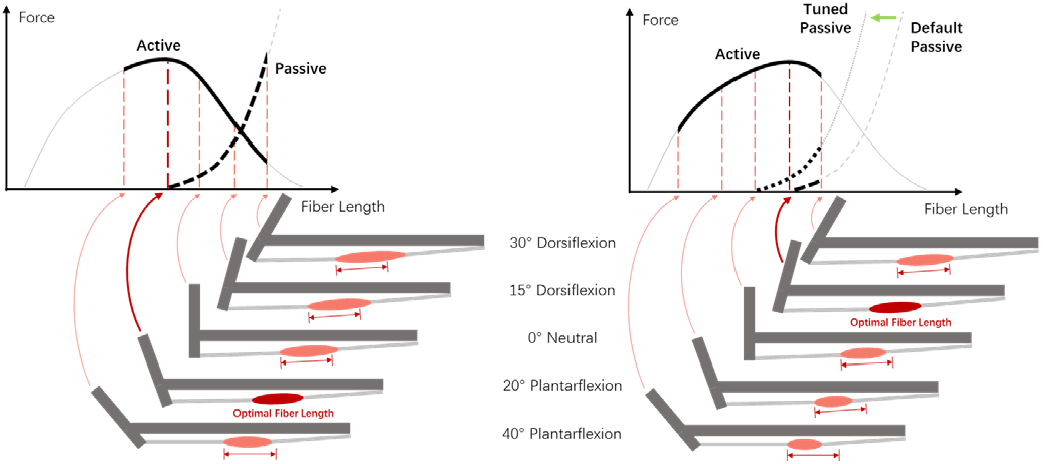
Optimal joint angle and *expressed* force–length curve. When undissected, there is a limit to which a muscle can be lengthened or shortened, depending on the joint’s range of motion (ROM). Thus, only part of the force– length curve can be expressed during movement. Left: If optimal fiber length is set at plantarflexion, the plantarflexors will be stretched as the ankle dorsiflexes to neutral and dorsiflexion positions, where active force capacity decreases and passive force dramatically increases. The right half of the active curve and a large portion of the passive curve are expressed. Right: If optimal fiber length is set at dorsiflexion, the plantarflexors will only stretched at large dorsiflexion angles. The left half of the active curve and only a small portion of the passive curve are expressed. Passive force can be increased by shifting the passive curve to the left.

It can be difficult to determine an optimal joint position. For example, Arnold et al. [25] first calculated the tendon slack length of the ankle muscles based on a resting ankle angle of 40° plantarflexion, as specified by Ward et al. [24]. However, the resultant tendon slack length values yielded excessive passive muscle forces, so tendon slack length was adjusted to a resting angle of 20° plantarflexion. This adjustment indicates that the resting ankle angle, where plantarflexion is large, is not appropriate for calculating tendon slack length. Ankle plantarflexors are short in plantarflexion positions and will be stretched to a large extent as the ankle dorsiflexes. If their optimality is set in high plantarflexion, then in dorsiflexion, along with the development of a very large passive force, the plantarflexors will perform on the descending limb of the active force–length curve (Fig. 3, left). From a physiological perspective, this is unlikely because plantarflexors are the main contributors in the push-off phase in a walking or running gait cycle, where a large moment of force is needed in dorsiflexion [62]–[64]. Maganaris [47], [65] measured plantarflexion moments independently contributed by the triceps surae and found them operating on the ascending limb of the active force– length curve with maximal force exerted in high dorsiflexion (Fig. 3, right).

In essence, the further away the fixation position is from the actual optimal position, the greater impact the resultant tendon slack length value will have on force estimation accuracy, as will be further demonstrated in the next section. Modelers and users should be exceptionally cautious about the joint position at which tendon slack length is set, and a reliable calculation requires experiment-based passive and active force–length relations. In Delp’s [21] lower limb model, the tendon slack length of each muscle was manually set for the total joint moments to peak at angles correspond to in vivo measurements. Simply speaking, *q*_*optz*_ is used instead of *q*_*meas*_.

A more straightforward approach is to directly measure and set tendon slack length [22], [23]. However, apart from having fiber optimality in the fixation position, this approach goes further and makes another simplification of *slack at rest*. A question mark remains as to whether tendons are slack when measured, since rigor mortis may fix cadavers in many poses passively unbalanced in vivo. Even if the values are accurate, they may not be directly applicable to the model, as musculoskeletal geometry need to match between the subject and the model for the measured tendon length to remain accurate as a modeling parameter.

We suggest that the concept of tendon slack length in modeling has little to do with *slackness* or *tendon*, but rather it acts like a phase-shift parameter defining the joint angle at which the fiber is at its optimality. It is neither conceptually biomechanical nor anatomical, so care should be taken to use measurements of tendon length without validation of kinetic data.

## IV. Sensitivity of Muscle Force Estimation

We have covered nine simplifications in the derivation of musculotendon parameters (Table II): The potential uncertainties are briefly discussed with examples given, but due to the nonlinear nature of muscle contraction dynamics, they are each differently associated with muscle force in contraction dynamics. Here in this section, we demonstrate to what extent errors in each musculotendon parameter influence force estimation and examine each simplification both numerically and analytically. This allows us to examine the effect of each simplification based a hypothetical range of errors to see how sensitive the model force estimation is to errors in the parameters.

**TABLE II.**
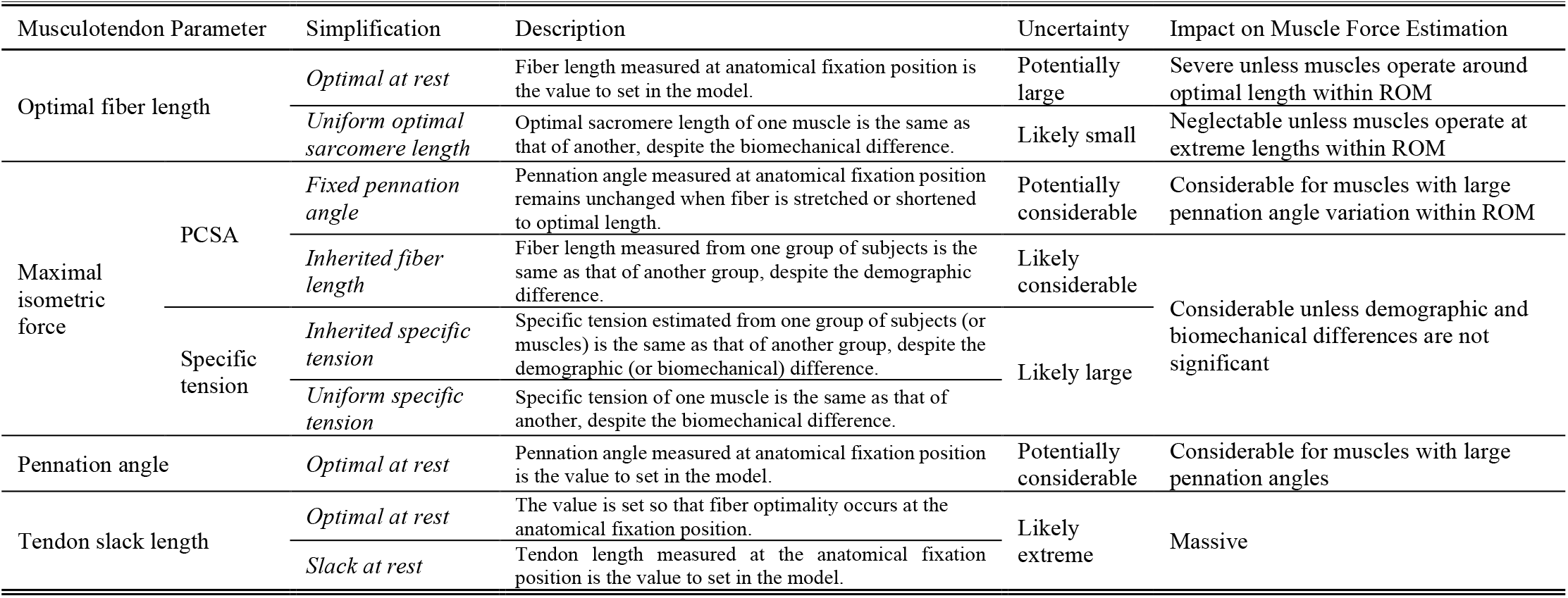
Simplifications in the derivation of musculotendon parameters

### A. Conventional Sensitivity Analysis

In order to perform the sensitivity analysis, a Hill-type muscle model was constructed based on Eqs. 1, 2, 3.1, 4.1, and 5 with force curve parameters from Hu et al. [11]. Isometric forces were calculated within the normalized fiber length range of 0.35–1.50 with maximal activation and a rigid tendon.

Fig. 4 shows the force–length curves when optimal fiber length, optimal pennation angle, and tendon slack length respectively changes, in comparison to ±10% and ±20% changes in PCSA or specific tension. If PCSA is independently obtained from optimal fiber length, then the impact is only noticeable at extreme lengths when the variation of optimal fiber length is greater than ±10% (Fig. 4, top left). However, if PCSA is derived with Eq. 3.1, the magnitude of force will change accordingly. Apart from the changes at extreme length, a ±10% variation of optimal fiber length is equivalent to a direct change of ±10% in maximal isometric force (Fig. 4, top right). Optimal pennation angle has a limited impact on force estimation, which is neglectable if both the actual and derived values are within 20°; the change is only large when pennation angle varies beyond 20° (Fig. 4, bottom left). The reason for this is simple: as the pennation angle increases, the small-angle approximation becomes invalid, and the increase of cosine value is no longer neglectable. As can be expected from the previous discussion, tendon slack length has the largest impact, where even an error as little as 1% would shift the curve for a force difference of nearly 10% (Fig. 4, bottom right).

**Fig. 4.**
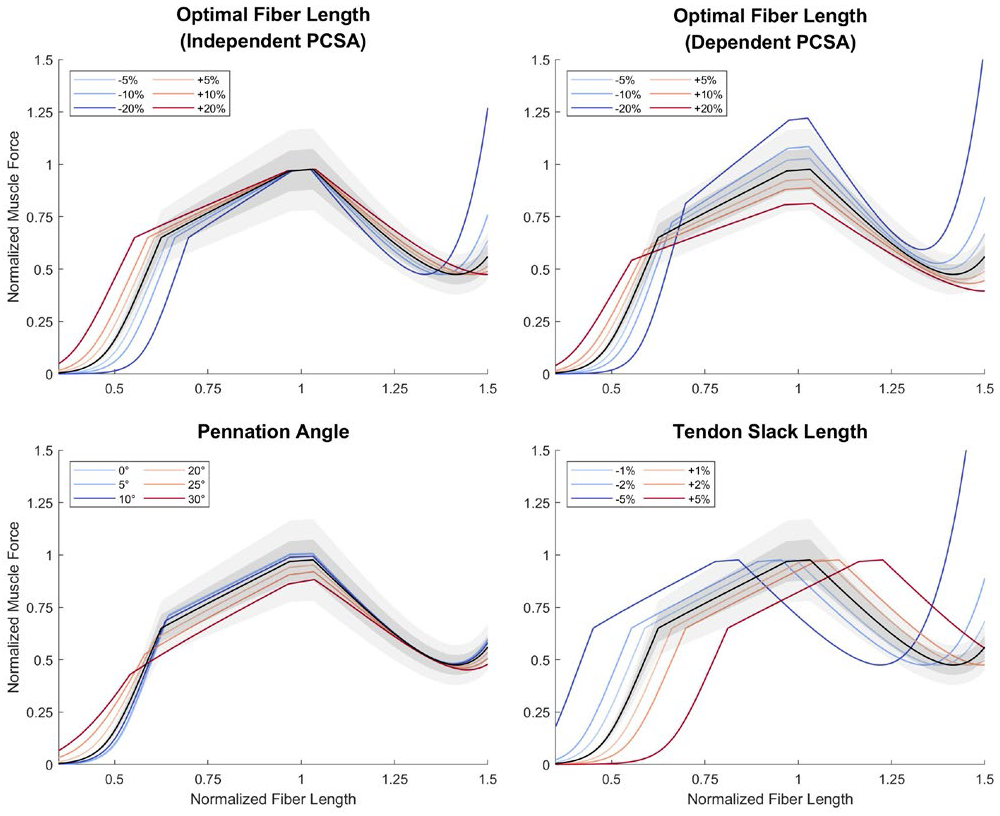
Normalized muscle force estimated with variations in musculotendon parameters in comparison with ±10% (dark gray) and ±20% (light gray) change in maximal isometric force. Top left: Muscle force estimated with variations in optimal fiber length (colored lines indicate amount of variation in parameters), when PCSA remains unchanged. Top right: Muscle force estimated with variations in optimal fiber length, when PCSA is calculated with Eq. 3.1. Bottom left: Muscle force estimated with variations in pennation angle. Bottom right: Muscle force estimated with variations in muscle length at optimal joint angle; referred to as tendon slack length for convenience. Note that each figure shows different ranges of parameter variation.

### B. Partial Derivatives of Muscle Force

To precisely compare the impact of each musculotendon parameter to force estimation, we derive the partial derivatives of the contraction dynamics in the above Hill-type muscle model.

Equations 6.1-6.4 are the partial derivatives of Eq. 1 with respect to optimal fiber length (the first equation considers independent PCSA, and the second has PCSA calculated using Eq. 3.1), optimal pennation angle, and MTU length at optimal joint position (denoted as *l*_*mt,opt*_), where:

- The original variable *q* in Eq. 1 is replaced by *l*_*mt*_ for convenience; in this case, *l*_*f,norm*_=(*l*_*mt*_-*l*_*t,sla*_)/(*l*_*f,opt*_ cos *α*) and *α* is calculated with Eq. 4.1. Musculoskeletal geometry can still be included if *l*_*mt*_(*q*) is modeled for substitution.
- *l*_*t,sla*_ is calculated using Eq. 5 with *l*_*mt,opt*_ as the first term.
- *f*_*ce*_ and *f*_*pe*_ are functions of *l*_*f,norm*_ respectively denoting the active and passive force–length curve.
- ∂*f*_*ce*_/∂*l*_*f,norm*_ and ∂*f*_*pe*_/∂*l*_*f,norm*_ are their partial derivatives, representing the gradients of the two curves.

These equations may also be conveniently implemented as optimization gradients for model calibration.

Fig. 5 shows the normalized values of the four partial derivatives. In general, a parameter can be regarded as more impactful to force estimation if the figure is more colorful, and trivial if it appears gray. The interpretation of the results is the same as the previous section, except that the partial derivatives offer a precise quantification of impact on force estimation from parameter variation.

**Fig. 5.**
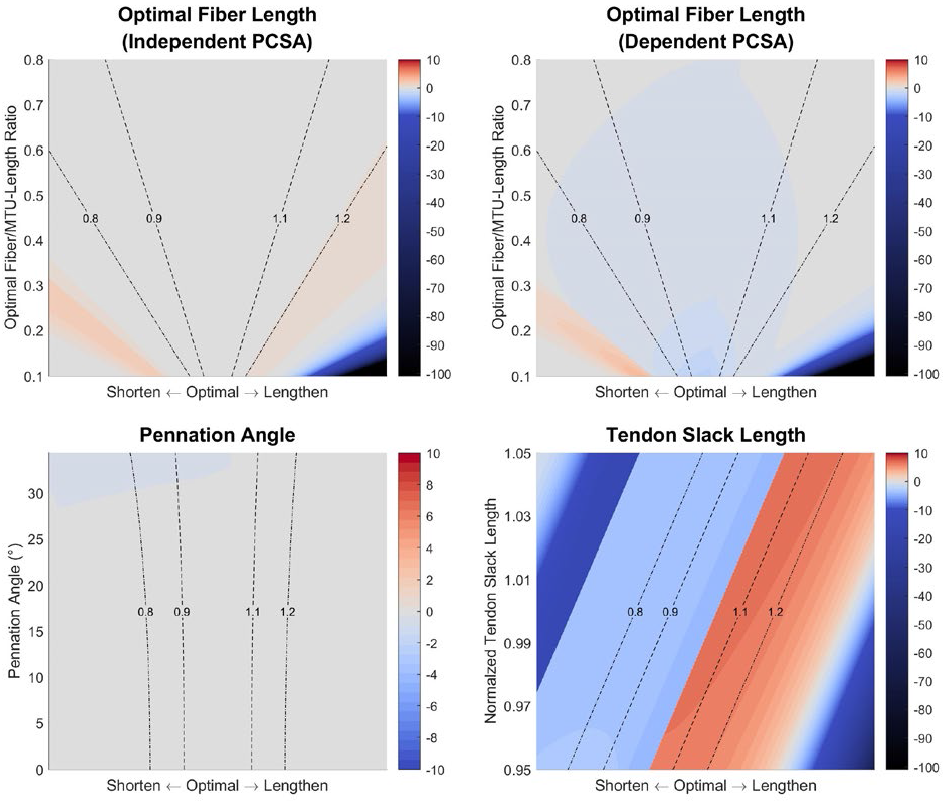
Normalized partial derivatives of muscle force with respect to musculotendon parameters. The value of the normalized partial derivative is indicated by the color on the heatmap: A darker red or blue indicates that, if the parameter is to increase from the given value indicated by the vertical axis, estimated muscle force is to increase or decrease by a larger extent, at the given muscle length indicated by the horizontal axis; the dashed lines are the contours of different normalized fiber lengths during shortening and lengthening. The color gray suggests the according change is small. For inter-parameter comparison, the value of the partial derivative is normalized by dividing the parameter value indicated by the vertical axis and maximal isometric force. Suppose the normalized partial derivative is 10, then if the parameter increases a sufficiently small portion (ε) of its original value, the estimated force capacity at the given muscle length will increase 10ε of its maximal isometric force. Top left: Normalized partial derivative with respect to optimal fiber length (expressed as in the ratio to the MTU length at optimal joint position), when PCSA remains unchanged. Top right: Normalized partial derivative with respect to optimal fiber length (expressed as in the ratio to the MTU length at optimal joint position), when PCSA is calculated with Eq. 3.1. Bottom left: Normalized partial derivative with respect to pennation angle. Bottom right: Normalized partial derivative with respect to muscle length at optimal joint angle (expressed as in the ratio to a constant value); referred to as tendon slack length for convenience.

### C. Inference on the Impact of Parameter Simplifications

#### Optimal fiber length

If PCSA is derived through calculation using optimal fiber length, then even a slight error in the latter will significantly change the force curve (Fig. 4, top right; Fig. 5, top right). Nevertheless, such an impact is essentially derived from the PCSA not from the optimal fiber length, so we hypothesize the scenario where PCSA is derived from alternative approaches irrelevant to other musculotendon parameters. A major problem in deriving optimal fiber length is when no sarcomere length values are available to perform optimal scaling, as is the case with Friederich and Brand (1990).

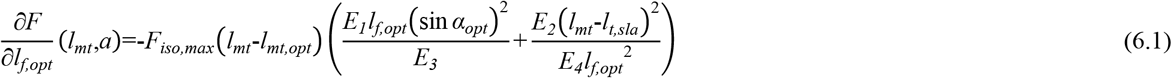

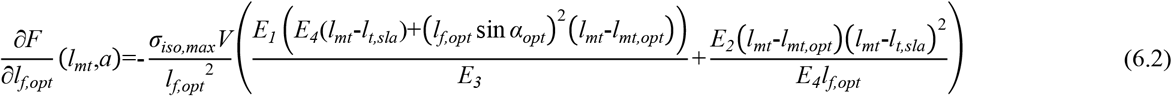

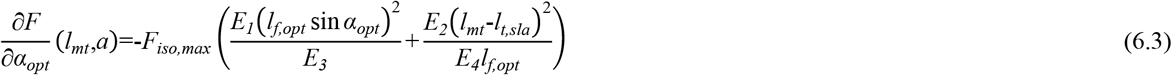

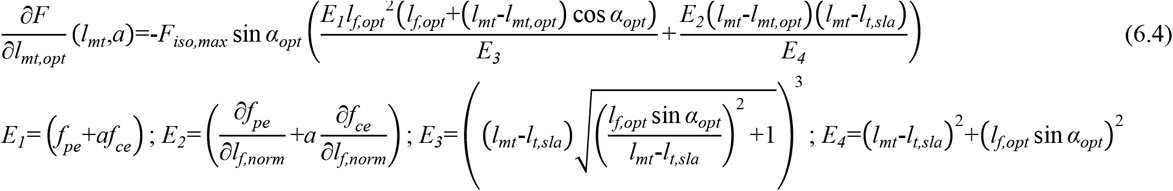

If we consider the situation in which they had measured the raw sarcomere lengths, and the values happen to be within the range of 2.2–3.2 μm, similar to the measurement of Ward et al. [24], then the difference between the unscaled and scaled values would be between 16–23%. For muscles that are fully shortened or lengthened within the range of motion (ROM), such errors might have the fiber overstretched at length where it should not be and vice versa (Fig. 4, top left), changing the expressed force curve. Next, suppose the true optimal fiber length is between 2.5–2.9 μm, then the error will be no more than 10% if raw fiber length is scaled by an optimal sarcomere length of 2.7 μm. In this case, the influence is only evident if the muscle operates at extreme lengths (Fig. 4, top left). Altogether, even if a uniform optimal sarcomere length is assumed, optimal scaling is still better than assuming optimal at rest.

#### Maximal Isometric Force

Force is constantly and evenly affected by errors in specific tension and PCSA (Eq. 2), the latter of which subjects to errors in muscle volume, optimal fiber length, and even pennation angle.

#### Pennation Angle

There is only noticeable change in force estimation if both pennation angle and the error are large enough (Fig. 4, bottom left; Fig. 5, bottom left). If Charles et al.’s [28] in vivo measurement is accurate, then all the lower limb muscles are free of such concern. Whereas according to Ward et al. [24], there are only four lower limb muscles with such large pennation angles as well as variances, namely the gluteus medius, glutes maximus, vasti medialis, and soleus. Yet they are some of the largest lower limb muscles whose absolute errors in simulation will be more significant. Caution should be given to these muscles if pennation angle is set as the value measured at the fixation position without correction.

#### Tendon slack length

The impact of an inaccurate tendon slack length is massive. The simplification of optimal at rest and slack at rest should be avoided, and if the parameter is set in this fashion without any kinetic calibration, simulation results will be far from satisfactory. Kinetic tuning, either manually [21] or automatically [61], is one practical approach; Equation 6.4 may be useful as optimization gradient.

## V. Summary

The biomechanical concept of *maximal force exertion* should always be borne in mind when dealing with musculotendon parameters. The current anatomical definitions can be misleading, and a more practical way is to redefine as follows:

- *Optimal fiber length*. The fiber length measured when muscle exerts active maximal isometric force. Alternatively, it can be defined as the fiber length scaled with the ratio of optimal to measured sarcomere length.
- *Optimal pennation angle*. The pennation angle measured when muscle exerts active maximal isometric force. Alternatively, it can be defined as the pennation angle scaled using nonlinear models such as Eqs. 4.1 and 4.2.
- *Optimal PCSA*. Calculated with muscle volume, optimal fiber length, and, if proven necessary, optimal pennation angle.
- *Optimal joint position*. The joint position where muscle exerts active maximal isometric force. This is the foundational parameter that cannot be calculated from muscle architecture data.

Many simplifications can be made when deriving musculotendon parameters, and depending on the muscle, they may negatively impact force estimation (Table II). We argue that automatically filling in each parameter with values from a dataset or an existing model can cause large uncertainties in the simulation results. Instead, the priority should be making sure that the parameters contain enough information about a muscle’s biomechanical properties, which is the exact issue with the simplifications in Table II.

To show that a set of musculotendon parameters is reasonably derived for modeling, modelers should specify how these typical simplifications are alternatively approached. In addition, if some parameters are inherited from previous work, then both the reasons behind this selection and the potential impact should be discussed. For example, if tendon slack length is manually tuned, the criterion to which each muscle is tuned must be stated: whether the resultant joint moments match with experimental data, or that certain joint angles are determined to be optimal. Or, if fiber length is derived from measured muscle length with the fiber- to muscle-length ratio of another subject group, explanation should be provided in case users are sensitive to certain demographic or anthropometric differences. Finally, we provide a flow chart that illustrates where each musculotendon parameter has been derived from for different models and datasets (Fig. 6). The goal is to provide a simple manner to see in what population certain parameters were determined from and how their values were derived. Both users and modelers could use this figure to check if the dataset or model they consider is free of concerning factors for their specific requirements. For instance, if surgeons wish to improve the prognosis of Achilles tendon rupture with information from a model simulation about the relation between fiber- to tendon-length ratio and heel raise height, then they first should select a model that properly sets tendon slack length. Fig. 6 would argue against using the three more recent models due to the simplification of either optimal at rest or slack at rest. Next, if it is shown that some of the optimal fiber length values are taken from Friederich and Brand [20] with the simplification of optimal at rest, then they should check that the soleus and gastrocnemius are not among the list. Then, seeing that pennation angle is taken from both datasets without optimal scaling, they might consider calibrating it with Eqs. 4.1 or 4.2 otherwise the large pennation angle of the soleus could induce error in force estimation. Finally, noticing that the subjects in the reference datasets are old and from a small sample, they should be careful about making deductions on young patients. Of course, some surgeons may believe that patient age plays a more important role in tendon biomechanics and thus prefer Rajagopal et al.’s [27] model. In this case, Fig. 6 would show that its fiber length is from elderly cadavers in Ward et al. [24], and in fact none of the current models have optimal fiber length derived from in vivo measurements on young subjects. After model selection, if the surgeons wishes to calibrate it with kinetic data collected from their patients, Eqs. 6.1-6.4 may be conveniently implemented as gradient in combination with the existing parameter optimization algorithm [61].

**Fig. 6.**
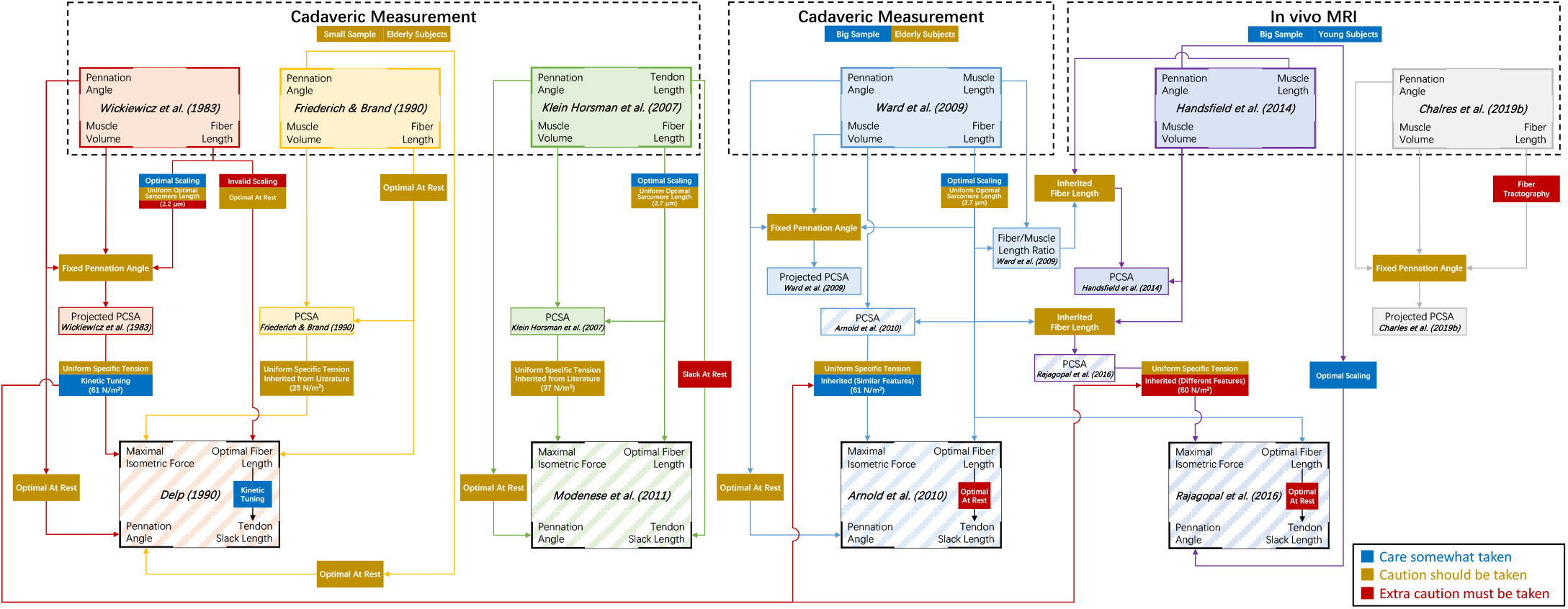
The derivation of musculotendon parameters in major lower limb muscle architecture datasets and models.

Despite the convenience, it is important to note that both Table II and Fig. 6 merely reflect potential uncertainties in contraction dynamics, and they should not be used to examine the overall simulation accuracy, which is a far more complicated concept. To give an example, in Eq. 1, the range of normalized fiber length is related to how muscle length changes within the ROM, which is neither determined by musculotendon parameters nor force curve parameters. This means that the expressed force–length curve may still be disfigured if muscle path is not accurately modeled. Moreover, even if the force curve is accurate, the estimated joint moment could still contain errors if the moment arm is inaccurate, which is no less problematic than inaccurate force estimation. Both muscle length and moment arm fall in the topic of musculoskeletal geometry, beyond the scope of this paper, but equally important.

## VI. Conclusion

As major factors in musculoskeletal modeling, musculotendon parameters can be overemphasized in model development by the recency of measurements, while having their derivation and impact on muscle force estimation overlooked. Here, we have outlined the derivation of all musculotendon parameters, highlighted simplifications in their estimates, and detailed the sensitivity of force estimation on these parameters.

Critically, musculotendon parameters represent specific biomechanical properties of the Hill-type model, therefore they cannot be fully calibrated by anatomical measurements alone. In particular, we highlight the case of tendon slack length, which, as a concept in modeling, has little to do with tendon in the anatomical sense. This makes tendon slack length difficult to measure or calculate. To calibrate musculotendon parameters with kinetic data such as joint moments, the partial derivatives of the Hill-type contraction dynamics are offered as gradients for fast and accurate parameter optimization.

In all, we provide information for the users of musculoskeletal models to decide the better-fitting datasets or models for their requirements in research or application. For the pursuit of extremely accurate kinetic estimation, we encourage the biomechanics community to focus on other model parameters (e.g., joint rotation center and muscle path), improve other model components (e.g., musculoskeletal geometry and neural control principle), and develop advanced methods for model calibration.

